# *De-novo* emergence and template switching of SINE retroposons during the early evolution of passerine birds

**DOI:** 10.1101/081950

**Authors:** Alexander Suh, Sandra Bachg, Stephen Donnellan, Leo Joseph, Jürgen Brosius, Jan Ole Kriegs, Jürgen Schmitz

**Affiliations:** Institute of Experimental Pathology (ZMBE), University of Münster, Münster, D-48149, Germany; Department of Evolutionary Biology (EBC), Uppsala University, Uppsala, SE-75236, Sweden; Evolutionary Biology Unit, South Australian Museum, Adelaide, SA 5000, Australia; Australian National Wildlife Collection, CSIRO National Research Collections Australia, Canberra, ACT 2601, Australia; Institute of Evolutionary and Medical Genomics, Brandenburg Medical School (MHB), Neuruppin, D-16816, Germany; LWL-Museum für Naturkunde, Westfälisches Landesmuseum mit Planetarium, Münster, D-48161, Germany

**Keywords:** transposon, retroposon, SINE, birds, Passeriformes, phylogenomics

## Abstract

Passeriformes (“perching birds” or passerines) make up more than half of all extant bird species. Here, we resolve their deep phylogenetic relationships using presence/absence patterns of short interspersed elements (SINEs), a group of retroposons which is abundant in mammalian genomes but considered largely inactive in avian genomes. The resultant retroposon-based phylogeny provides a powerful and independent corroboration of previous indications derived from sequence-based analyses. Notably, SINE activity began in the common ancestor of Eupasseres (passerines excl. the New Zealand wrens Acanthisittidae) and ceased before the rapid diversification of oscine passerines (songbirds). Furthermore, we find evidence for very recent SINE activity within suboscine passerines, following the emergence of a SINE via acquisition of a different tRNA head as we suggest through template switching. We propose that the early evolution of passerines was unusual among birds in that it was accompanied by activity of SINEs. Their genomic and transcriptomic impact warrants further study in the light of the massive diversification of passerines.

## Introduction

Short interspersed elements (SINEs) are the most abundant group of the reverse-transcribed retroposons in mammalian genomes (Sotero-Caio, et al. in revision). They rely on *trans*-mobilization by the enzymatic machinery of long interspersed elements (LINEs) (Ohshima, et al. 1996), a *parasitic* interaction so successful that the human genome contains >1,500,000 SINEs compared to <900,000 LINEs (Lander, et al. 2001). On the other hand, SINEs are scarce in avian genomes, which has been noted as one of the most peculiar genomic features of birds (Hillier, et al. 2004; Warren, et al. 2010; Zhang, et al. 2014). While LINEs exhibit up to 700,000 copies in avian genomes, there are only 6,000-17,000 SINEs per avian genome (Zhang, et al. 2014), most of these ancient and heavily degraded (Kapusta and Suh 2016). Presence/absence patterns of SINEs in orthologous genomic loci are rare genomic changes appreciated widely as virtually homoplasy-free phylogenetic markers (Shedlock, et al. 2004; Ray, et al. 2006). Given the aforementioned scarcity of SINEs, it is not surprising that the emergence and activity of SINEs has never been studied in birds. On the other hand, other types of retroposed elements (REs; LINEs from the chicken repeat 1 superfamily, CR1, and long terminal repeat elements, LTRs) have helped resolve the relationships of various groups of birds, such as Galliformes (Kaiser, et al. 2007; Kriegs, et al. 2007; Liu, et al. 2012), Neoaves (Suh, Paus, et al. 2011; Matzke, et al. 2012; Suh, Smeds, et al. 2015), Palaeognathae (Haddrath and Baker 2012; Baker, et al. 2014), and others (St. John, et al. 2005; Watanabe, et al. 2006; Suh, et al. 2012; Kuramoto, et al. 2015). In the meantime, the sequencing of dozens of avian genomes has revealed SINEs with putative lineage specificity (Warren, et al. 2010; Kapusta and Suh 2016; Suh, et al. 2016) and thus the potential for conducting presence/absence analyses in specific groups of birds.

Here we conduct, to our knowledge, the first study of the emergence and activity of SINEs in birds. We focus on the deep phylogenetic relationships of passerines, the largest radiation of birds with nearly 6,000 extant species (Barker, et al. 2004), using 44 presence/absence markers of SINEs and other REs. In contrast to the only previous study of retroposons in passerines with a single RE marker (Treplin and Tiedemann 2007), our multilocus dataset permits the reassessment of sequence-based phylogenies [e.g., (Barker, et al. 2004; Selvatti, et al. 2015; Moyle, et al. 2016)] and, simultaneously, the reconstruction of the temporal activity of SINEs and other REs during early passerine evolution.

## Results and Discussion

We initially chose RE marker candidates from selected retroposon families of zebra finch [including TguSINE1, (Warren, et al. 2010)] in October 2009, a time when genome assemblies were available only for chicken and zebra finch (Hillier, et al. 2004; Warren, et al. 2010). Candidates for presence/absence loci were therefore identified via pairwise alignment of RE-flanking sequences from zebra finch to orthologous regions in chicken (Materials and Methods). This was followed by *in-vitro* presence/absence screening of RE marker candidates as detailed elsewhere (Suh, Kriegs, et al. 2011; Suh, Paus, et al. 2011) using a representative taxon sampling of all major groups of passerines *sensu* Barker, et al. (2004) (Supplementary Table S1). We complemented this with a screening of GenBank (http://www.ncbi.nlm.nih.gov/genbank/) for additional SINEs, which identified a TguSINE1-like insertion in *myoglobin* intron 2 of *Pitta anerythra* (accession number DQ785977) that is absent in the orthologous position of other *Pitta* species (Irestedt, et al. 2006). We termed this element “PittSINE” and identified PittSINE marker candidates in a DNA sample of *Pitta sordida* via inter-SINE PCR [(Kaukinen and Varvio 1992); Materials and Methods]. This was followed by cloning and sequencing of the 500-bp to 1,000-bp fraction of PCR amplicons, alignment to chicken and zebra finch genomes to reconstruct the left and right SINE-flanking regions, and then *in-vitro* presence/absence screening of PittSINE marker candidates.

Next, we characterized the structural organization of passerine SINEs (Fig. 1) using the available TguSINE1 consensus sequence (Warren, et al. 2010) and after generating a majority-rule consensus of PittSINE insertions in our sequenced presence/absence markers (Supplementary Data S1). Both SINEs have highly similar, CR1-derived tails (Fig. 1) which exhibit the typical hairpin for putative binding by the CR1 reverse transcriptase and an 8-bp microsatellite at their very end for target-primed reverse transcription (Suh 2015). However, the heads of these SINEs are derived from different tRNA genes, namely tRNA^Ile^ in TguSINE1 and tRNA^Asp^ in PittSINE (Fig. 1). Sequence alignment suggests that the tRNA-derived SINEs heads are more similar to the respective tRNA genes than they are to each other (Fig. 1C). However, the opposite is the case for the CR1-derived SINE tails, which exhibit four diagnostic nucleotides distinguishing them from the highly similar 3’ end of CR1-X1_Pass (Fig. 1C).

**Figure 1:**
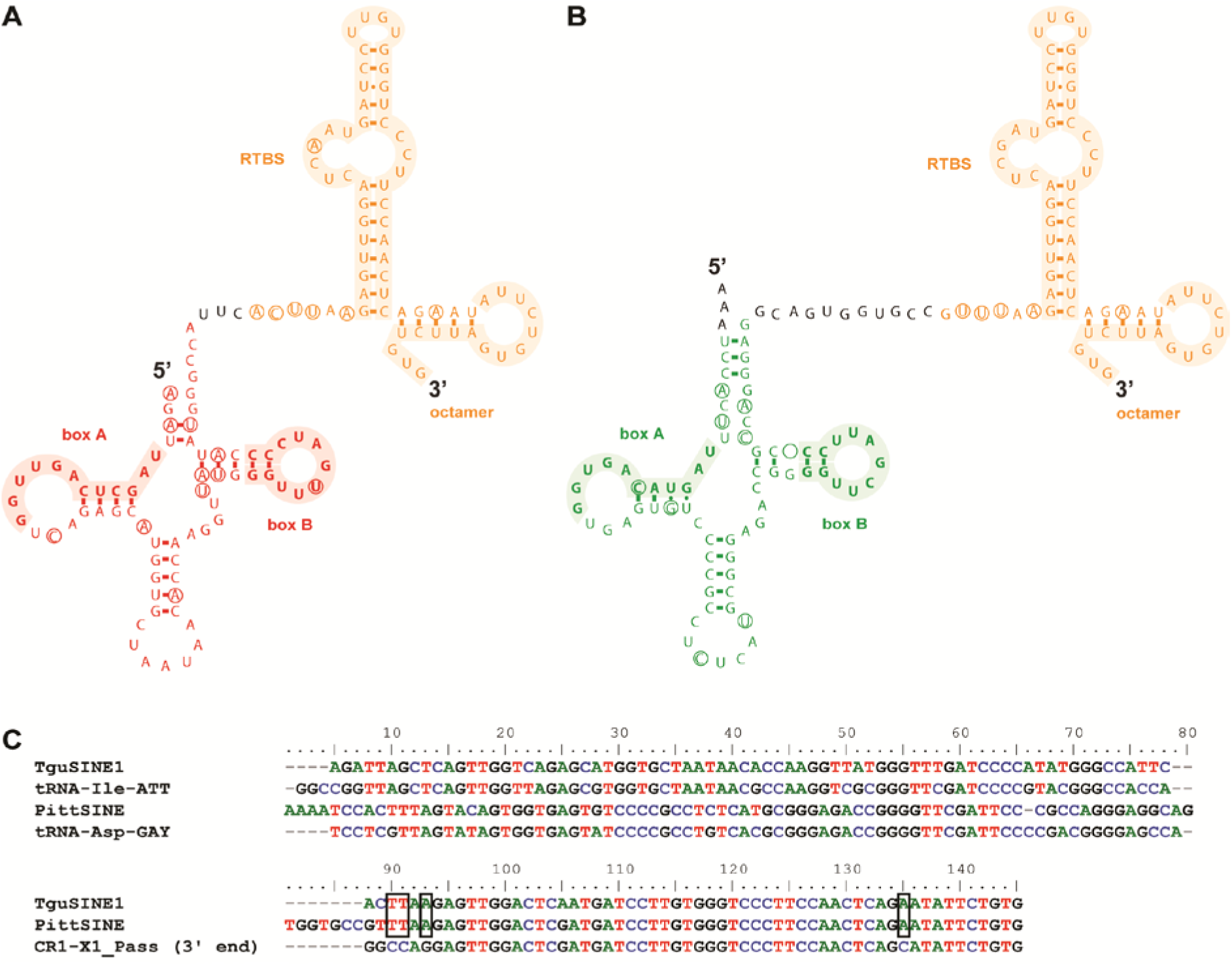
Simplified RNA secondary structures of passerine SINEs with CR1-derived tails (orange) and tRNA-derived heads. The SINE heads are tRNA^Ile^ (red) in TguSINE1 (**A**) and tRNA^Asp^ (green) in PittSINE (**B**). Shaded regions denote promoter boxes A and B in tRNAs, as well as the reverse transcriptase binding site (RTBS) and 5’-ATTGTRTG-3’ microsatellite typical for CR1 elements of amniotes (Suh 2015). Circles indicate nucleotide differences between SINE consensus sequences and the respective tRNAs or CR1 they are derived from. The RTBS hairpin structure is also visible in mfold-based predictions of SINE secondary structure (Supplementary Fig. S1). (**C**) DNA alignment of TguSINE1 and PittSINE with respective tRNA genes and 3’ end of CR1-X1_Pass. Black boxes denote diagnostic nucleotides present in the CR1-derived tails of TguSINE1 and PittSINE.

We further investigated this peculiar pattern using phylogenetic analyses of the CR1-derived SINE tails and avian CR1 subfamilies *sensu* Suh, Churakov, et al. (2015), which again suggests that TguSINE1 and PittSINE have a single SINE ancestor which derived its tail from CR1-X1_Pass (Fig. 2A). Assuming that SINEs are *trans*-mobilized by LINE reverse transcriptase enzymes due to high sequence similarity between SINE tails and LINE 3’ ends (Ohshima, et al. 1996) and thus depend on LINE activity, the most likely candidate for SINE mobilization is the CR1-X1_Pass subfamily. This is further supported by temporal overlap of TguSINE1 and CR1-X activity in RE landscapes of the zebra finch genome (Fig. 2B). Additionally, we detected direct evidence for temporal overlap of TguSINE1 and CR1-X1_Pass activity through our presence/absence analyses (Fig. 3A, Supplementary Table S1).

**Figure 2:**
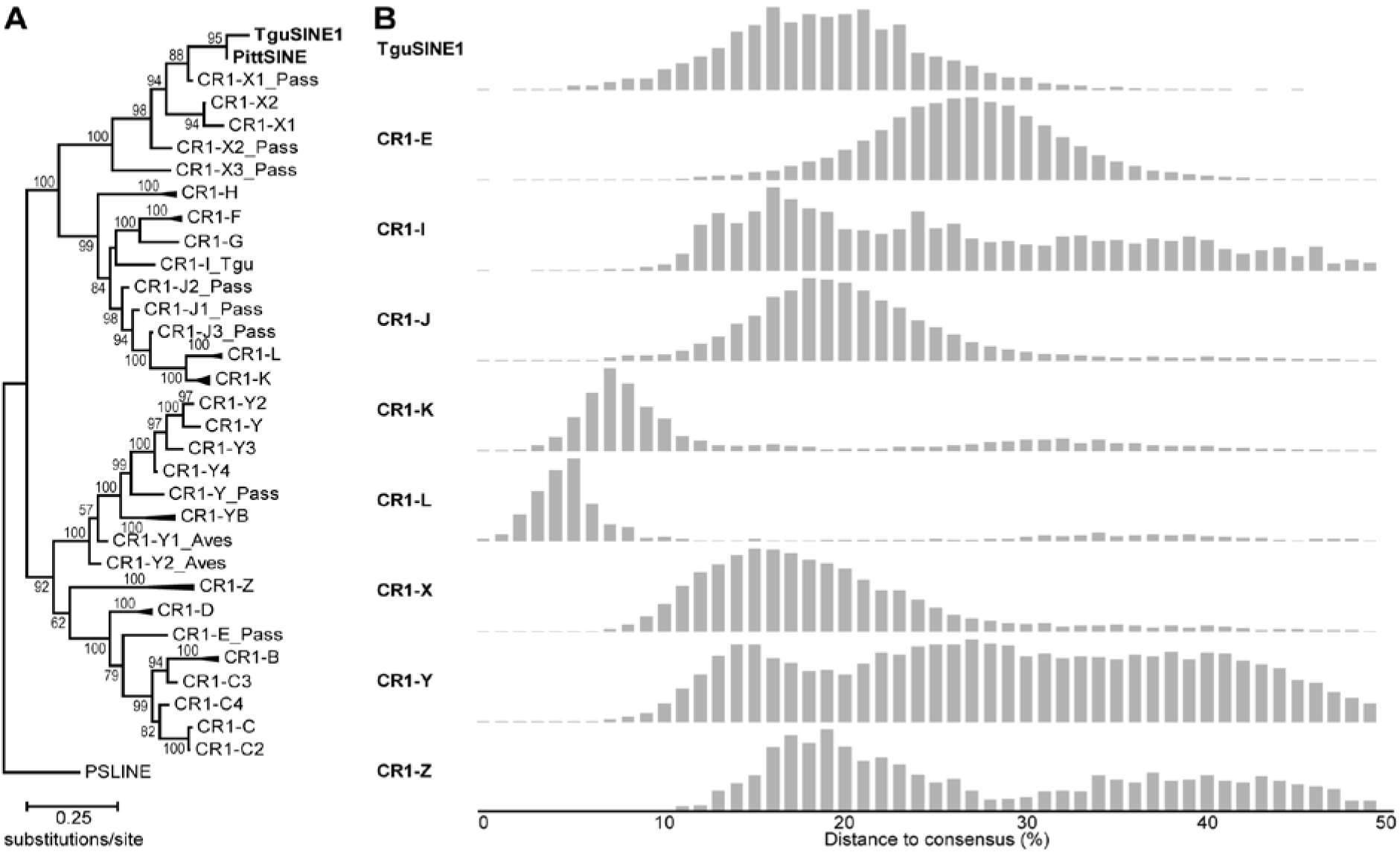
Passerine SINEs share a common ancestor and are mobilized by CR1-X.xs. (**A**) Maximum likelihood phylogeny of passerine SINE tails and avian CR1 subfamilies in Repbase (Jurka, et al. 2005) (GTRCAT model, 1,000 bootstrap replicates) suggests that TguSINE1 and PittSINE arose from the same CR1-X subfamily (CR1-X1_Pass) and share a common SINE ancestor. Note that the topology of the CR1 phylogeny is identical to that of previous studies (Suh, et al. 2012; Suh, Churakov, et al. 2015). (**B**) Comparison of the TguSINE1 landscape with landscapes of CR1 families (merged subfamilies from panel **A**) suggests temporal overlap of SINE and CR1-X activity in the zebra finch genome. RE landscapes were generated using the zebra finch assembly taeGut2 following methods detailed elsewhere (Suh, Churakov, et al. 2015).

**Figure 3:**
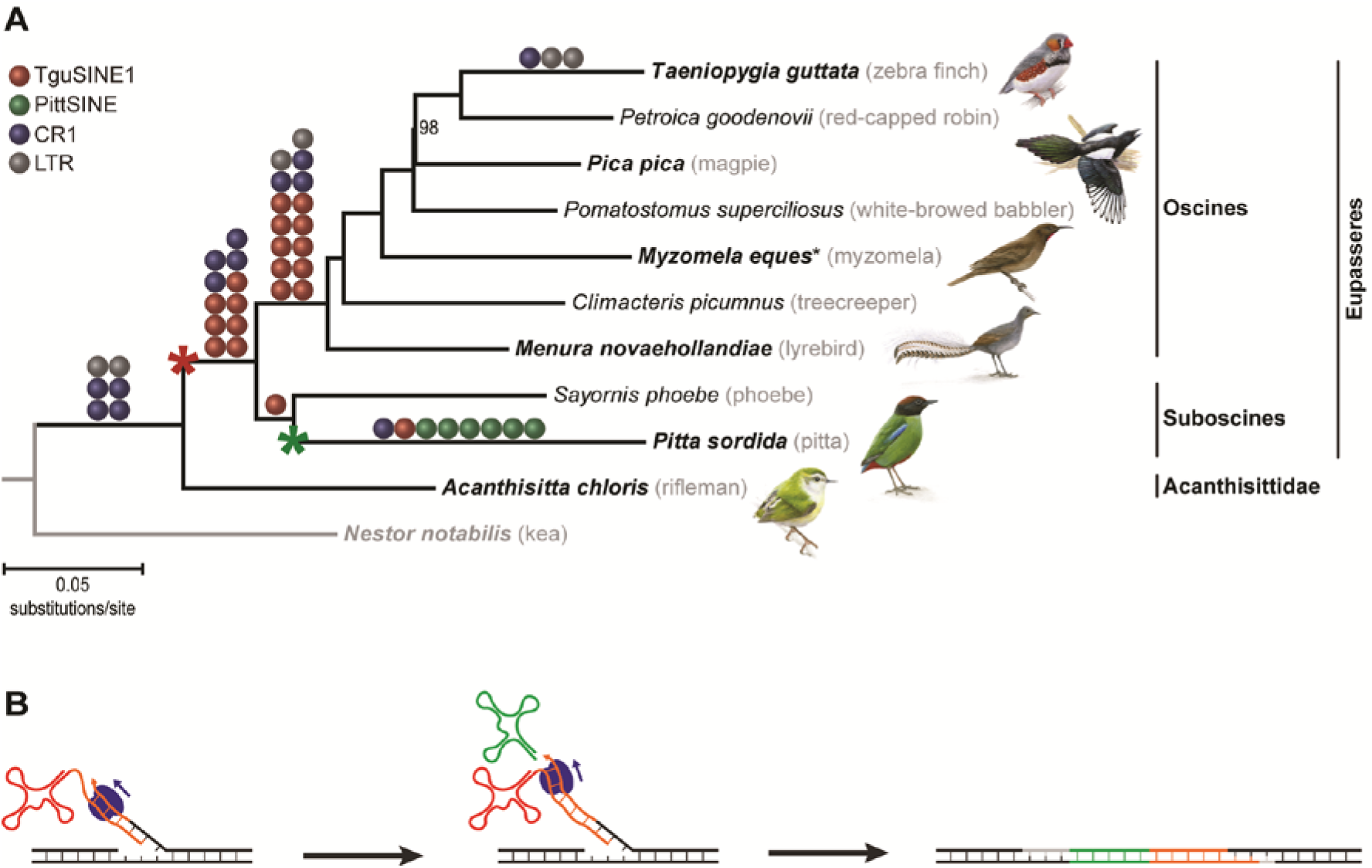
Emergence and timing of CR1-mobilized SINE activity during early passerine evolution. (**A**) Phylogenomic analysis of early passerine relationships using retroposon presence/absence markers (colored balls) mapped on a maximum likelihood phylogeny of concatenated retroposon-flanking sequences (GTRCAT model, 1,000 bootstrap replicates). Our sampling consists of the major deep passerine lineages *sensu* Barker et al. (2004). Red and green asterisks indicate emergence of TguSINE1 and PittSINE, respectively. The black asterisk indicates that for some loci (Supplementary Table S1), *Malurus cyaneus* was sampled instead of *Myzomela eques* to represent the Maluridae/Meliphagidae clade (Barker, et al. 2004). Only bootstrap values <100% are shown and the names of pictured birds are emphasized in bold. (**B**) A scenario for the emergence of PittSINE. Template switching from TguSINE1 RNA (red, tRNA^Ile^ head; orange, CR1 tail) to tRNA^Asp^ (green) during target-primed reverse transcription by CR1 reverse transcriptase (blue). The resultant tRNA^Asp^-CR1 chimaera was flanked by a target site duplication (grey) and transcriptional activation gave rise to the PittSINE family.

Our extensive RE presence/absence analyses yielded 19 TguSINE1 markers, 6 PittSINE markers, 13 CR1 markers, and 6 LTR markers which we could trace across a representative taxon sampling of the major groups of passerines *sensu* Barker, et al. (2004). Careful inspection of presence/absence alignments using strict criteria (see Materials and Methods) yielded a conflict-free set of RE markers, which we mapped on a maximum likelihood tree constructed from concatenated RE-flanking sequences from the same data set (Fig. 3A). For three of the deepest passerine branching events, we found a multitude of RE markers and thus statistically significant support in available RE marker tests (Waddell, et al. 2001; Kuritzin, et al. 2016). These relationships are the respective monophyly of passerines and oscines, as well as the monophyly of Eupasseres (Mayr and Manegold 2004), a group comprising all passerines except the New Zealand wrens Acanthisittidae. The Eupasseres/Acanthisittidae split was first observed in sequence analyses of few nuclear genes (Barker, et al. 2002; Ericson, et al. 2002) and has since been recovered in ever-growing nuclear sequence analyses [e.g., (Barker, et al. 2004; Ericson, et al. 2014; Selvatti, et al. 2015; Moyle, et al. 2016)]. Our analysis of rare genomic changes thus provides the first assessment of this group using an independent marker type and phylogenetic method. None of our RE markers inserted during the rapid radiation of oscine passerines, however, sequence analysis of the RE-flanking regions yielded a topology identical to the aforementioned previous studies. Of particular interest are the four deep-branching oscine lineages Menuridae (e.g., *Menura novaehollandiae*), Climacteridae (e.g., *Climacteris picumnus*), Maluridae/Meliphagidae (e.g., *Malurus cyaneus* and *Myzomela eques*), and Pomatostomidae (e.g., *Pomatostomus superciliosus*) because these four lineages together have been rarely included in passerine phylogenetic studies. We find a branching order (Fig. 3A) which recapitulates previous phylogenetic estimates based on few nuclear genes (Barker, et al. 2004) or ultraconserved elements (Moyle, et al. 2016). This suggests that the rapid radiation of oscines can be congruently resolved even with non-genome-scale data. We note that this is in contrast to the neoavian radiation, which appears to be partially irresolvable even with retroposon markers [reviewed by Suh (2016)]. Within passerines, we further note that the conflict between single-RE support for a Picathartidae/Corvidae clade (Treplin and Tiedemann 2007) and sequence-based phylogenies (Han, et al. 2011) results from incorrect placing of this RE marker on the passerine Tree of Life (see legend of Supplementary Fig. S2 for more information).

We then traced the emergence and activity of SINEs across the passerine Tree of Life. Given that RE marker candidates were initially chosen on chicken/zebra finch alignments, we expect no bias in the distribution of RE markers on the lineage leading to zebra finch. TguSINE1 was mostly active in the ancestor of oscines and, to a lesser extent, in the ancestor of Eupasseres. Interestingly, we find no evidence for TguSINE1 activity in the common ancestor of passerines or during/after the radiation of oscines and therefore hypothesize that TguSINE1 emerged in Eupasseres and became extinct in the oscines ancestor (Fig. 3A). The emergence of TguSINE1 is thus the first “genome morphology” character for the monophyly of Eupasseres and supplements support from skeletal morphology, which is limited to the presence of a ‘six-canal pattern’ in the hypotarsus (Manegold, et al. 2004).

In contrast to the situation in oscines, the activity of TguSINE1 appears to have been longer in suboscines, postdating the divergence between Old World and New World suboscines (i.e., pitta and phoebe in Fig. 3A). This recent, potentially lineage-specific activity coincides with the putative restriction of PittSINEs to Old World suboscines (e.g., *Pitta* spp.). The aforementioned indication for a common SINE ancestor of TguSINE1 and PittSINE evidenced by four diagnostic nucleotides in their CR1-derived SINE tails (cf. Fig. 1C and Fig. 2A) suggests that the younger PittSINE emerged from the older TguSINE1 after acquisition of a new tRNA-derived head. Assuming that TguSINE1 and PittSINE were both active on the pitta lineage, we propose that the most plausible mechanism for PittSINE emergence was template switching from TguSINE1 to a nearby tRNA during reverse transcription (Fig. 3B). Template switching has been previously proposed in a wide range of chimeric retroposons [e.g., (Brosius 1999; Gilbert and Labuda 2000; Buzdin, et al. 2002; Nishihara, et al. 2016)] and appears to be a particularly common opportunity for SINEs to *parasitize* different LINEs via acquisition of new SINE tails (Ohshima and Okada 2005; Nishihara, et al. 2016). Our data show that template switching may also happen for SINE heads and we speculate that the acquisition of a new SINE head from a different tRNA gene may provide intact and active promoter components for efficient transcription by RNA polymerase III.

## Conclusions

Here, we reconstructed the deep phylogenetic relationships of passerines using presence/absence patterns of unusual SINE insertions and other REs. This permitted us to follow the emergence, activity, and extinction of TguSINE1 and PittSINE across the evolution of the most species-rich group of birds. While this SINE activity was considerably lower than, for example, that in mammals, it nevertheless exemplifies that at least some birds have a more diverse repetitive element landscape than previously anticipated. Furthermore, we note that the activity of TguSINE1 appears to coincide with the evolution of vocal learning during early passerine evolution (Suh, Paus, et al. 2011). Previous evidence suggests that ~4% of birdsong-associated transcripts in the zebra finch brain contain retroposons (Warren, et al. 2010) and it thus remains to be seen whether SINE activity influenced the evolution of, for example, vocal learning in oscine passerines.

## Materials and Methods

We identified candidates for presence/absence loci for TguSINE1 and other selected zebra finch retroposons via pairwise alignment of RE loci from zebra finch to orthologous regions in chicken. This was done by comparing and extracting the respective RE-flanking sequences in the UCSC Genome Browser (Fujita, et al. 2011), followed by automatic alignment using MAFFT version 6 (Katoh and Toh 2008). In order to find PittSINE marker candidates, we conducted inter-SINE PCR (Kaukinen and Varvio 1992) using a single, PittSINE-specific oligonucleotide primer (5’-CTCGTTAGTATAGTGGTGAGTGTC-3’) and standard PCR parameters of Suh, Kriegs, et al. (2011) with 50°C annealing temperature. All presence/absence screenings were done using oligonucleotide primers binding to conserved RE-flanking regions in chicken/zebra finch alignments (Supplementary Table S2), using the touchdown PCR and cloning protocols of Suh, Paus, et al. (2011).

For each presence/absence marker candidate, we first aligned all sequences automatically using MAFFT (E-INS-I option) and then manually inspected these for misalignments. We considered a marker candidate as phylogenetically informative and reliable “*if, in all species sharing this RE, it featured an identical orthologous genomic insertion point (target site), identical RE orientation, identical RE subtype, identical target site duplications (direct repeats, if present) and a clear absence in other species*” (Suh, Paus, et al. 2011). This led to a total of 44 high-quality RE presence/absence markers (Supplementary Table S1, Supplementary Data S2). All newly generated sequences were deposited in GenBank (accession numbers XXXXXXXX-XXXXXXXX).

All maximum likelihood sequence analyses were conducted using RAxML 8.1.11 (Stamatakis, et al. 2008) on the CIPRES Science Gateway (Miller, et al. 2010). For the CR1 phylogeny, we used the alignment from Suh et al. (2012), excluded grebe-specific CR1 elements, and added the CR1-derived tails of TguSINE1 and PittSINE. For the passerine phylogeny, we removed the RE sequences from our presence/absence alignments and concatenated the remaining RE-flanking sequences into a multilocus alignment (Supplementary Data S3).

## Abbreviations

CR1: chicken repeat 1.
LINE: long interspersed element.
Mb: million basepairs.
MY: million years.
MYA: million years ago.
RE: retroposed element.
RT: reverse transcriptase.
RTBS: reverse transcriptase binding site.
SINE: short interspersed element.

## Acknowledgments

We thank Tim Pock, Meike Hüdig, and Felix Babatz for help with *in-vitro* experiments, and Gerald Mayr and Gennady Churakov for helpful discussions. We are grateful to Leanne Wheaton, Simone Schehka (Allwetterzoo Münster), Robert Palmer, Stephanie Hodges, Geoffrey E. Hill, Franziska A. Franke, Sharon Birks (Burke Museum), and Werner Beckmann (LWL-DNA-und Gewebearchiv) for providing blood and tissue samples, and to Jón Baldur Hlíðberg for generating the bird paintings. This research was funded by the Deutsche Forschungsgemeinschaft (KR3639 to J.O.K. and J.S.).

## Supplementary Material

**Supplementary Figure S1:**
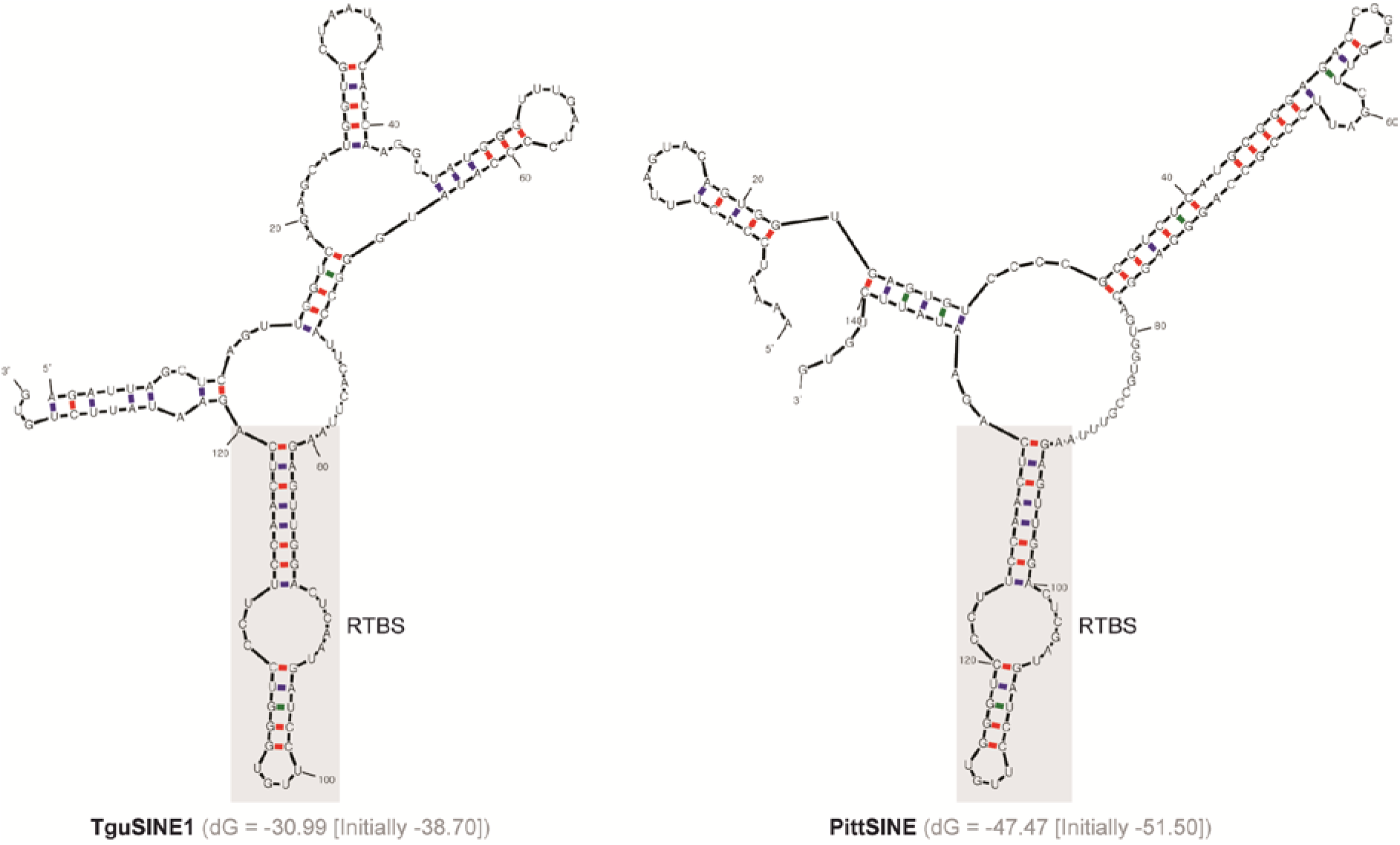
Predictions of secondary structures of TguSINE1 and PittSINE using mfold (Zuker 2003).

**Supplementary Figure S2:**
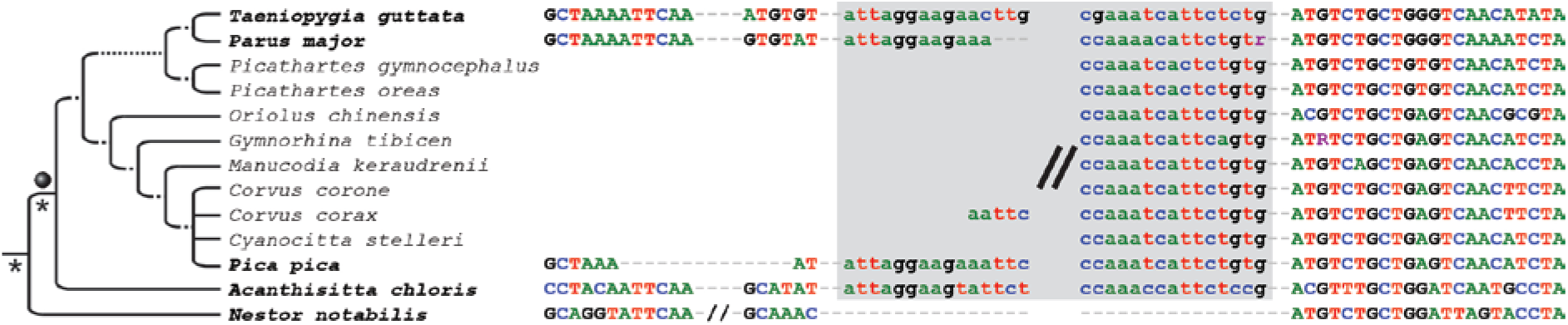
The RE marker of Treplin & Tiedemann (2007) does not suggest “*phylogenetic affinity of rockfowls (genus* Picathartes*) to crows and ravens (Corvidae)*”. Our extended phylogenetic sampling instead suggests that the RE insertion occurred in the ancestor of all passerines (grey ball). This discrepancy is because Treplin & Tiedemann (2007) were, due to methodological limitations, unable to detect RE presence in non-oscine passerines (*Acanthisitta chloris*) and RE absence in the parrot outgroup (*Nestor notabilis*). Taxa with bold names were sampled in the present study and the grey box denotes the 5’ and 3’ end of the CR1 insertion.

**Supplementary Table S1:**
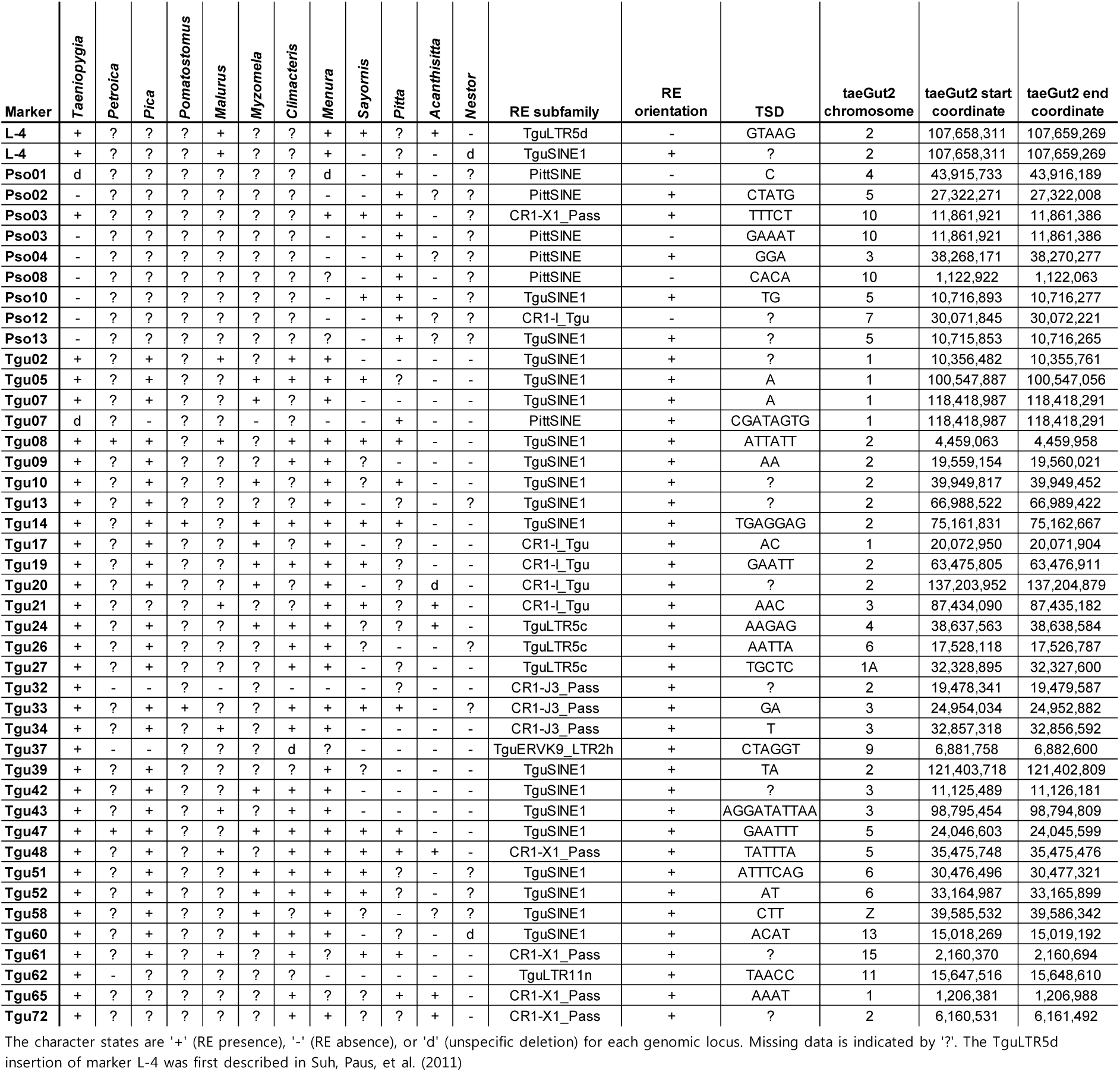
Presence/absence matrix of passerine retroposon markers including RE target site duplication (TSD) motifs and location in the taeGut2 assembly of the zebra finch genome.

**Supplementary Table S2:**
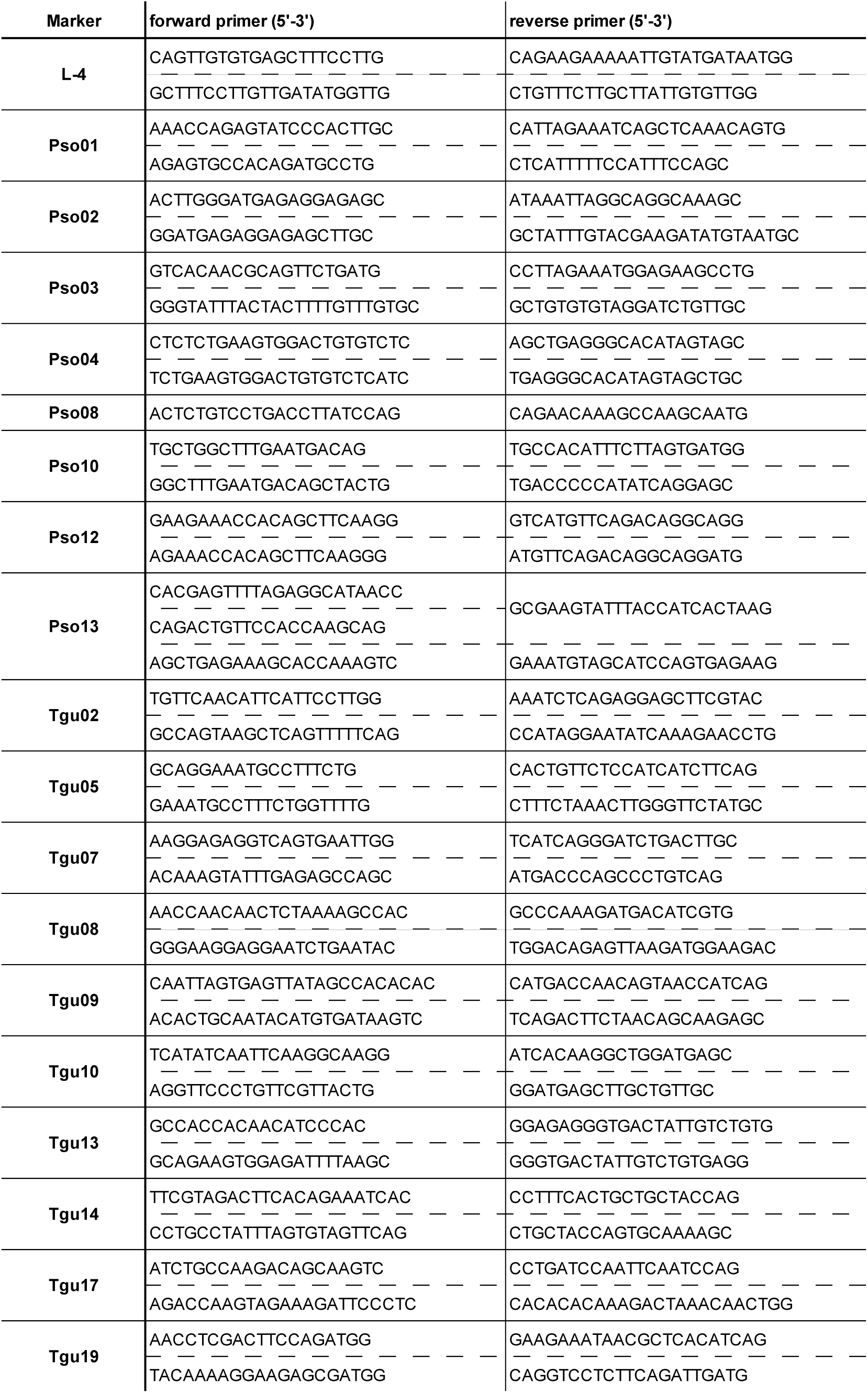
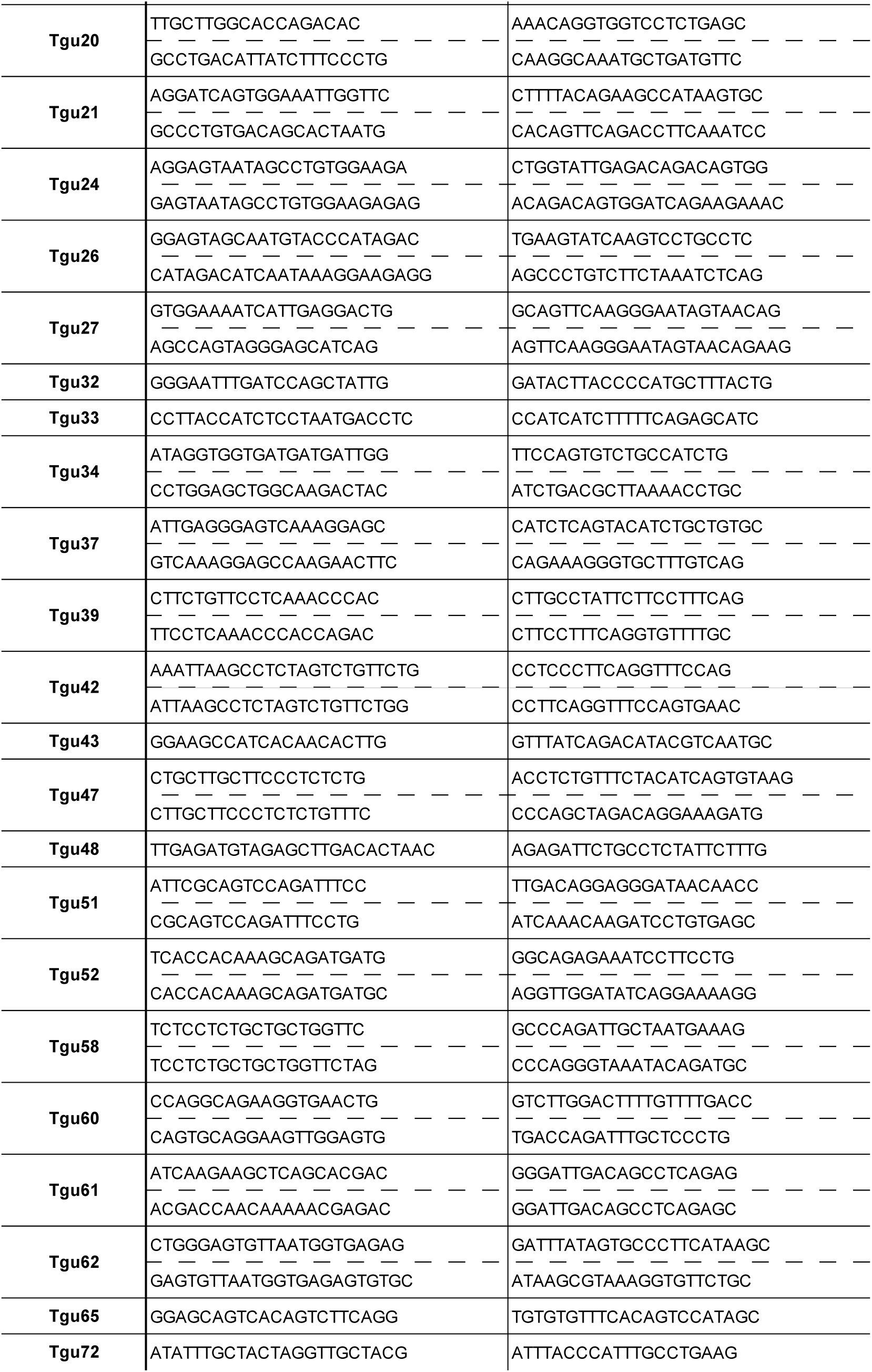
Oligonucleotide primer sequences.

**Supplementary Data S1: Majority-rule consensus sequence for PittSINE as reconstructed from our PittSINE-bearing presence/absence patterns.**

**Supplementary Data S2: Fasta-formatted alignments of all RE presence/absence markers.**

**Supplementary Data S3: Fasta-formatted multilocus alignment of concatenated RE-flanking sequences used for generating the phylogenetic tree of Fig. 3A**.

## References

Baker AJ, Haddrath O, McPherson JD, Cloutier A. 2014. Genomic support for a moa-tinamou clade and adaptive morphological convergence in flightless ratites. Mol Biol Evol 31:1686–1696.

Barker FK, Barrowclough GF, Groth JG. 2002. A phylogenetic hypothesis for passerine birds: taxonomic and biogeographic implications of an analysis of nuclear DNA sequence data. Proceedings of the Royal Society of London Series B: Biological Sciences 269:295–308.

Barker FK, Cibois A, Schikler P, Feinstein J, Cracraft J. 2004. Phylogeny and diversification of the largest avian radiation. Proc Natl Acad Sci USA 101:11040–11045.

Brosius J. 1999. Genomes were forged by massive bombardments with retroelements and retrosequences. Genetica 107:209–238.

Buzdin A, Ustyugova S, Gogvadze E, Vinogradova T, Lebedev Y, Sverdlov E. 2002. A new family of chimeric retrotranscripts formed by a full copy of U6 small nuclear RNA fused to the 3' terminus of L1. Genomics 80:402–406.

Ericson P, Klopfstein S, Irestedt M, Nguyen J, Nylander J. 2014. Dating the diversification of the major lineages of Passeriformes (Aves). BMC Evol Biol 14:8.

Ericson PG, Christidis L, Cooper A, Irestedt M, Jackson J, Johansson US, Norman JA. 2002. A Gondwanan origin of passerine birds supported by DNA sequences of the endemic New Zealand wrens. Proc R Soc B 269:235–241.

Fujita P, Rhead B, Zweig A, Hinrichs A, Karolchik D, Cline MS, Goldman M, Barber G, Clawson H, Coelho A DM, Dreszer TR, Giardine BM, Harte RA, Hillman-Jackson J, Hsu F, Kirkup V, Kuhn RM, Learned K, Li CH, Meyer LR, Pohl A, Raney BJ, Rosenbloom KR, Smith KE, Haussler D, Kent WJ. 2011. The UCSC Genome Browser database: update 2011. Nucleic Acids Res 39:D876–D882.

Gilbert N, Labuda D. 2000. Evolutionary inventions and continuity of CORE-SINEs in mammals. J Mol Biol 298:365–377.

Haddrath O, Baker AJ. 2012. Multiple nuclear genes and retroposons support vicariance and dispersal of the palaeognaths, and an Early Cretaceous origin of modern birds. Proceedings of the Royal Society B 279:4617–4625.

Han K-L, Braun EL, Kimball RT, Reddy S, Bowie RCK, Braun MJ, Chojnowski JL, Hackett SJ, Harshman J, Huddleston CJ, et al. 2011. Are transposable element insertions homoplasy free?: an examination using the avian tree of life. Syst Biol 60:375–386.

Hillier LW, Miller W, Birney E, Warren W, Hardison RC, Ponting CP, Bork P, Burt DW, Groenen MA, Delany ME, et al. 2004. Sequence and comparative analysis of the chicken genome provide unique perspectives on vertebrate evolution. Nature 432:695716.

Irestedt M, Ohlson JI, Zuccon D, Källersjö M, Ericson PGP. 2006. Nuclear DNA from old collections of avian study skins reveals the evolutionary history of the Old World suboscines (Aves, Passeriformes). Zool Scripta 35:567–580.

Jurka J, Kapitonov VV, Pavlicek A, Klonowski P, Kohany O, Walichiewicz J. 2005. Repbase Update, a database of eukaryotic repetitive elements. Cytogenet Genome Res 110:462467.

Kaiser VB, van Tuinen M, Ellegren H. 2007. Insertion events of CR1 retrotransposable elements elucidate the phylogenetic branching order in galliform birds. Mol Biol Evol 24:338–347.

Kapusta A, Suh A. 2016. Evolution of bird genomes -a transposon's-eye view. Ann N Y Acad Sci in press.

Katoh K, Toh H. 2008. Recent developments in the MAFFT multiple sequence alignment program. Brief Bioinform 9:286–298.

Kaukinen J, Varvio S-L. 1992. Artiodactyl retroposons: association with microsatellites and use in SINEmorph detection by PCR. Nucleic Acids Res 20:2955–2958.

Kriegs JO, Matzke A, Churakov G, Kuritzin A, Mayr G, Brosius J, Schmitz J. 2007. Waves of genomic hitchhikers shed light on the evolution of gamebirds (Aves: Galliformes). BMC Evol Biol 7:190.

Kuramoto T, Nishihara H, Watanabe M, Okada N. 2015. Determining the position of storks on the phylogenetic tree of waterbirds by retroposon insertion analysis. Genome Biol Evol 7:3180–3189.

Kuritzin A, Kischka T, Schmitz J, Churakov G. 2016. Incomplete lineage sorting and hybridization statistics for large-scale retroposon insertion data. PLoS Comput Biol 12:e1004812.

Lander ES, Linton LM, Birren B, Nusbaum C, Zody MC, Baldwin J, Devon K, Dewar K, Doyle M, Fitzhugh W, et al. 2001. Initial sequencing and analysis of the human genome. Nature 409:860–921.

Liu Z, He L, Yuan H, Yue B, Li J. 2012. CR1 retroposons provide a new insight into the phylogeny of Phasianidae species (Aves: Galliformes). Gene 502:125–132.

Manegold A, Mayr G, Mourer-Chauviré C. 2004. Miocene songbirds and the composition of the European passeriform avifauna. Auk 121:1155–1160.

Matzke A, Churakov G, Berkes P, Arms EM, Kelsey D, Brosius J, Kriegs JO, Schmitz J. 2012. Retroposon insertion patterns of neoavian birds: strong evidence for an extensive incomplete lineage sorting era. Mol Biol Evol 29:1497–1501.

Mayr G, Manegold A. 2004. The oldest European fossil songbird from the early Oligocene of Germany. Naturwissenschaften 91:173–177.

Miller MA, Pfeiffer W, Schwartz T editors. Proceedings of the Gateway Computing Environments Workshop (GCE). 2010 2010: New Orleans, LA.

Moyle RG, Oliveros CH, Andersen MJ, Hosner PA, Benz BW, Manthey JD, Travers SL, Brown RM, Faircloth BC. 2016. Tectonic collision and uplift of Wallacea triggered the global songbird radiation. Nat Commun 7:12709.

Nishihara H, Plazzi F, Passamonti M, Okada N. 2016. MetaSINEs: Broad distribution of a novel SINE superfamily in animals. Genome Biol Evol 8:528–539.

Ohshima K, Hamada M, Terai Y, Okada N. 1996. The 3' ends of tRNA-derived short interspersed repetitive elements are derived from the 3' ends of long interspersed repetitive elements. Molecular and Cellular Biology 16:3756–3764.

Ohshima K, Okada N. 2005. SINEs and LINEs: symbionts of eukaryotic genomes with a common tail. Cytogenet Genome Res 110:475–490.

Ray DA, Xing J, Salem A-H, Batzer MA. 2006. SINEs of a nearly perfect character. Syst Biol 55:928–935.

Selvatti AP, Gonzaga LP, Russo CAdM. 2015. A Paleogene origin for crown passerines and the diversification of the Oscines in the New World. Mol Phylogenet Evol 88:1–15.

Shedlock AM, Takahashi K, Okada N. 2004. SINEs of speciation: tracking lineages with retroposons. Trends Ecol Evol 19:545–553.

Sotero-Caio C, Platt II RN, Suh A, Ray DA. in revision. Evolution and diversity of transposable elements in vertebrates. Genome Biol Evol.

St. John J, Cotter J-P, Quinn TW. 2005. A recent chicken repeat 1 retrotransposition confirms the Coscoroba-Cape Barren goose clade. Mol Phylogenet Evol 37:83–90.

Stamatakis A, Hoover P, Rougemont J. 2008. A rapid bootstrap algorithm for the RAxML web servers. Syst Biol 75:758–771.

Suh A. 2016. The phylogenomic forest of bird trees contains a hard polytomy at the root of Neoaves. Zool Scripta 45:50–62.

Suh A. 2015. The specific requirements for CR1 retrotransposition explain the scarcity of retrogenes in birds. J Mol Evol 81:18–20.

Suh A, Churakov G, Ramakodi MP, Platt II RN, Jurka J, Kojima KK, Caballero J, Smit A, Vliet K, Hoffmann FG, et al. 2015. Multiple lineages of ancient CR1 retroposons shaped the early genome evolution of amniotes. Genome Biol Evol 7:205–217.

Suh A, Kriegs JO, Brosius J, Schmitz J. 2011. Retroposon insertions and the chronology of avian sex chromosome evolution. Mol Biol Evol 28:2993–2997.

Suh A, Kriegs JO, Donnellan S, Brosius J, Schmitz J. 2012. A universal method for the study of CR1 retroposons in nonmodel bird genomes. Mol Biol Evol 29:2899–2903.

Suh A, Paus M, Kiefmann M, Churakov G, Franke FA, Brosius J, Kriegs JO, Schmitz J. 2011. Mesozoic retroposons reveal parrots as the closest living relatives of passerine birds. Nat Commun 2:443.

Suh A, Smeds L, Ellegren H. 2015. The dynamics of incomplete lineage sorting across the ancient adaptive radiation of neoavian birds. PLoS Biol 13:e1002224.

Suh A, Witt CC, Menger J, Sadanandan KR, Podsiadlowski L, Gerth M, Weigert A, McGuire JA, Mudge J, Edwards SV, et al. 2016. Ancient horizontal transfers of retrotransposons between birds and ancestors of human pathogenic nematodes. Nat Commun 7:11396.

Treplin S, Tiedemann R. 2007. Specific chicken repeat 1 (CR1) retrotransposon insertion suggests phylogenetic affinity of rockfowls (genus Picathartes) to crows and ravens (Corvidae). Mol Phylogenet Evol 43:328–337.

Waddell PJ, Kishino H, Ota R. 2001. A phylogenetic foundation for comparative mammalian genomics. Genome Informatics 12:141–154.

Warren WC, Clayton DF, Ellegren H, Arnold AP, Hillier LW, Kϋnstner A, Searle S, White S, Vilella AJ, Fairley S, et al. 2010. The genome of a songbird. Nature 464:757–762.

Watanabe M, Nikaido M, Tsuda TT, Inoko H, Mindell DP, Murata K, Okada N. 2006. The rise and fall of the CR1 subfamily in the lineage leading to penguins. Gene 365:57–66.

Zhang G, Li C, Li Q, Li B, Larkin DM, Lee C, Storz JF, Antunes A, Greenwold MJ, Meredith RW, et al. 2014. Comparative genomics reveals insights into avian genome evolution and adaptation. Science 346:1311–1320.

Zuker M. 2003. Mfold web server for nucleic acid folding and hybridization prediction. Nucleic Acids Res 31:3406–3415.

